# The role of familiarity in grouping decisions of Trinidadian guppies

**DOI:** 10.64898/2026.11.06.764979

**Authors:** Adithi S. Rao, Nina P. C. Alves, Jon T. Sakata, Simon M. Reader

## Abstract

Social grouping is widely observed in numerous taxa, providing individuals with a variety of benefits including efficient information exchange and predator protection. However, grouping can also carry costs such as increased competition for resources, which means that individuals must often make decisions regarding joining or leaving a group. These decisions to leave or stay might in turn depend on prior experience with potential groupmates. Past work has shown that many fish distinguish between familiar and unfamiliar conspecifics and use these distinctions in their grouping choices. Trinidadian guppies (*Poecilia reticulata*), well known for their diversity in shoaling preferences, are an excellent system in which to investigate the role of familiarity in grouping decisions. We examined preferences for familiar individuals following both a 15-day and 55-day cohabitation period in two different guppy populations. Utilizing two binary-choice designs, we found no evidence for a familiarity preference in either population, despite observing a high preference to group with conspecifics in both populations. These results suggest that various contextual- and population-level differences may impact familiarity preferences, in turn contributing to the growing body of work on the evolution of social organization and grouping decisions in teleost species and beyond.

## Introduction

Social grouping is a widespread and important phenomenon across the animal kingdom (Krause and Ruxton 2002). Grouping can carry costs, such as increased competition and disease transmission (Wilson et al. 2003; Schmid-Hempel 2021), but can also confer numerous benefits such as foraging opportunities, access to social information, and reduced predation risk (Neill and Cullen 1974; Mooring and Hart 1992; Barber and Wright 2001; Ward and Webster 2016). These costs and benefits mean that individuals often make active choices about both when and with whom to group, such as grouping with individuals of the same species, population or sex (Wiley 2013; Ward and Webster 2016). Numerous cues can be used to discriminate between individuals, including visual, behavioural, auditory and chemosensory cues, either alone or in combination (Toccalino et al. 2016; Sayigh et al. 1999; Karavanich and Atema 1998; de la Zerda et al. 2022; Tibbetts and Dale 2007). Importantly, these cues may be learned, as in the case of familiarity: the recognition of other individuals based on prior social experience (Griffiths 2003; Ward et al. 2020).

Indeed, grouping advantages have been shown to be particularly pronounced when individuals associate with familiar individuals. In multiple taxa, familiarity can improve group cohesion, buffer stress, and reduce intraspecific conflict (Denommé and Mason, 2022; Silk 2007; Reddon et al. 2021; Ward et al. 2020). Familiarity can also provide a mechanism of kin recognition (Waldman 1987; Ward & Hart 2003). Grouping is well studied in teleost fish, with numerous species making decisions on whether and with whom to group (Krause and Godin 1994; Griffiths and Magurran 1998; Krause and Ruxton 2002; Hoare and Krause 2003). Familiarity specifically is also well studied, and associating with familiar conspecifics is known to further enhance anti-predator responses such as detection, cohesion, and coordination, as well as the learning of foraging routes (Chivers et al. 1995; Barber and Wright 2001; Nadler et al. 2021; Swaney et al. 2001; Lucon-Xiccato et al. 2022).

The freshwater guppy, *Poecilia reticulata*, typically forms shoals (Magurran 2005), and female guppies use familiarity to assess potential shoaling partners, for example preferring to group with individuals they have previously been housed with (Griffiths and Magurran 1997a; Lachlan et al. 1998). Griffiths and Magurran (1997a) found that female guppies associated preferentially with familiar conspecifics (their own tank-mates) only after a period of 12 days of cohabitation. This preference remained even following a 5-week period of separation from each other (Bhat and Magurran 2006). Further, shoaling preferences for familiar individuals diminished as the number of individuals living together increased, suggesting the fish were learning individuals (Griffiths and Magurran 1997b). Given the described benefits of familiarity, it is important to establish the conditions under which familiarity preferences are expressed and whether previous findings are replicated in different settings.

While replication and reproducibility are widely acknowledged as essential to scientific progress, animal studies can present particular challenges for reproducibility since local conditions can interact with genetic background to impact trait development and expression (Webster and Rutz 2020; von Kortzfleisch et al. 2022). It is thus important to establish to what extent results generalize across animals with different backgrounds and local conditions (Webster and Rutz 2020; von Kortzfleisch et al. 2022). Here, we adopted previous methods (Griffiths and Magurran 1997a) to examine familiarity preferences in two populations of female guppies following both a 15-day and 55-day cohabitation period using two different experimental apparatuses. This experiment consists of three parts (hereafter ‘Tests 1-3’). In Test 1, we examined preferences for familiar conspecifics in the two different guppy populations. Subjects of the same populations were housed in groups of six that were classified as either ‘familiar’ (conspecifics from the same housing tank) or ‘unfamiliar’ (conspecifics from any other housing tank). Following 15 days of cohabitation with conspecifics, we gave subjects a binary-choice preference test wherein they could pick between a shoal consisting of familiar or unfamiliar individuals (Test 1). Fifteen days has previously been shown to be sufficient for familiarity to develop in female guppies (Griffiths and Magurran 1997a). Based on our negative results from Test 1, we tested the same subjects in a modified testing set-up (Test 2) following 40 additional days of cohabitation. After Test 2, we performed a preference test for a shoal versus an empty tank (Test 3) to verify that our procedure and apparatus could detect any shoaling preferences, should they exist. We predicted that as in Griffiths and Magurran (1997a), subjects would demonstrate a preference for shoals and specifically for familiar individuals.

## Test 1

### Materials and Methods

Subjects and stimulus fish were adult female guppies from two laboratory-maintained populations and had not been previously used in other studies. The ‘Aripo/Quare-cross’ population are descendants resulting from a cross between two high-predation populations collected in the Lower Aripo and Quare rivers in Trinidad in 2009 and 2010 (Gotanda 2015). The ‘mixed wild origin’ population are of mixed Trinidadian origin, a gift of the Rodd Laboratory (University of Toronto, Canada) in 2018. Both populations have since been maintained as outbred laboratory populations. Before the experiment, fish were housed in 150L mixed-sex stock tanks of approximately 50-70 individuals each. In previous work, similar group sizes were large enough to prevent familiarity development among individuals (Griffiths and Magurran 1997b). Stock tanks included filters, gravel substrate, a heater, artificial aquarium plants, and a terracotta pot for enrichment, and were maintained at 25°C and a 12:12 light-dark cycle. Weekly 30% water changes and water testing (pH, hardness, nitrites, nitrates, and ammonia) were performed on stock tanks. Fish were fed daily with tropical fish flakes (TetraMin, Tetra, Germany), supplemented with decapsulated brine shrimp eggs (*Artemia sp.*, Brine Shrimp Direct, USA) three times a week.

We chose 60 female fish haphazardly from their stock home tanks, avoiding visibly gravid individuals, and grouped them in tanks (*l* 40 x *w* 20 x *h* 25 cm) of 6 fish from the same population, visually and olfactorily isolated from other tanks. While stock tanks contained a mix of kin and non-kin, the haphazard selection meant that kinship was randomized across housing groups. On the morning of the test, 3 fish in each tank were randomly chosen to be subjects and the remaining 3 were designated as stimulus fish. The body lengths of the subjects (mean ± SE = 19.84 ± 0.394 mm) and the stimulus fish (20.58 ± 0.903 mm) were not significantly different from each other (Welch two-sample t-test: t(19.5) = 0.755, p = 0.46). Individuals were first tested after being housed in these tanks for a period of 15 days with identical feeding and housing conditions as the stock tanks, including enrichment. Fish that shared a single tank were considered ‘familiar’, while fish in different tanks were considered ‘unfamiliar’. Testing was performed during daylight hours beginning at approximately 8:00 daily. Fish were fed on their normal diet and schedule after testing finished for the day and were not food-deprived prior to testing.

### Test 1: Familiarity Preference

We conducted a preference test modelled on Griffiths and Magurran (1997a) (Figure 1). Stimulus shoal fish from the same population were placed 6 cm from either end of the testing tank (*l* 90 x *w* 45 x *h* 30 cm; water depth 20 cm) in two perforated and clear 1 litre plastic bottles (diameter: 8 cm) with the tops cut off and covered with plastic cups. Perforations (about 4 mm in diameter, 5 cm apart, around the circumference of the bottle) allowed chemosensory cues to pass without allowing physical contact between subjects and stimulus fish. A shoal of 3 fish was placed in each bottle, with familiar fish on one side (the 3 non-subjects from the same tank as the subject), and unfamiliar fish on the other (3 non-subject fish chosen randomly from any other one tank, ensuring each tank’s stimulus fish were picked an equal number of times overall). An individual subject (n=30) was then moved by net into a glass cylinder in the centre of the testing tank to acclimate for 15 minutes. When released, the subject could thus shoal with the familiar or unfamiliar fish (or remain outside the shoaling zones). Video was recorded from above for 15 minutes using a Sony 4K Camcorder (Handycam FDR-AX100).

**Figure 1:**
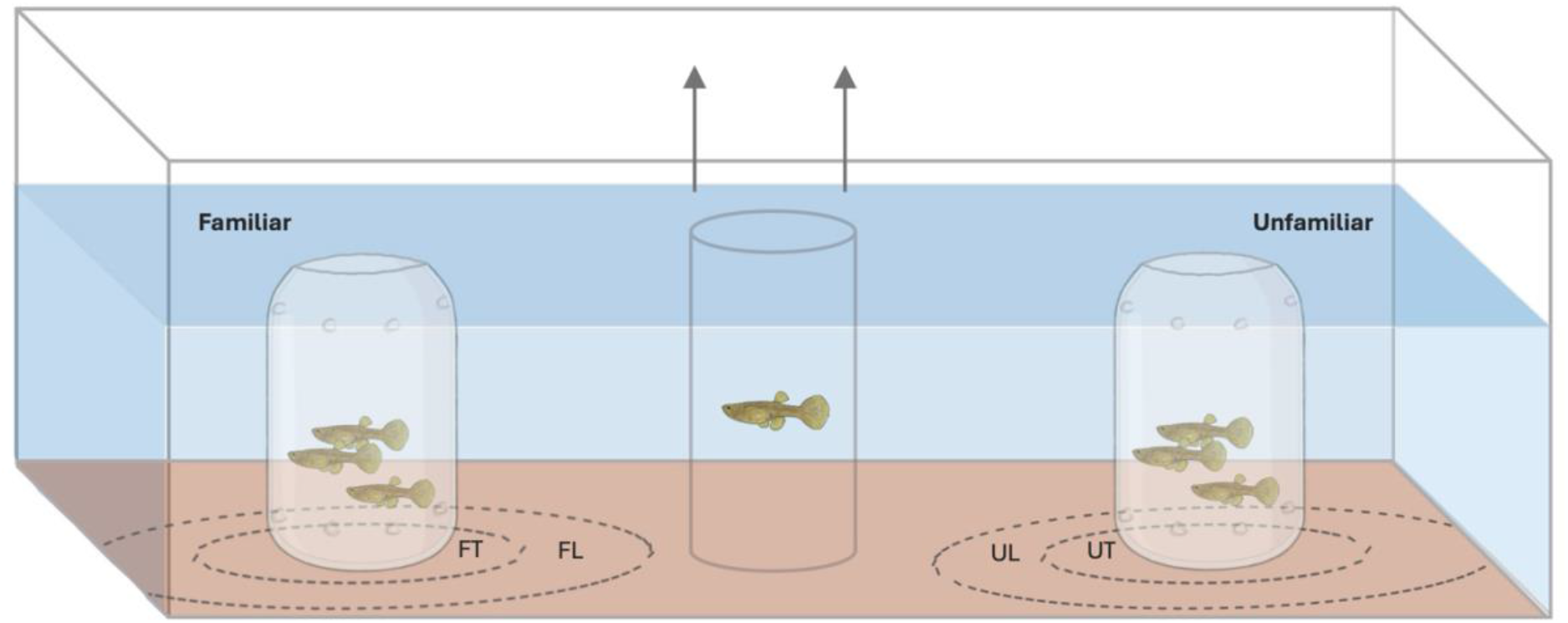
Test 1 familiarity preference test set-up, front view. Subject is released from a central cylinder and allowed to swim freely between a familiar and unfamiliar shoal of three fish, each contained in clear perforated plastic bottles. Fish are illustrated larger than actual size, image not to scale. Shoaling zones were defined as 0-5 and 0-10 cm from each shoal: FT - familiar tight shoaling zone; FL - familiar loose shoaling zone; UT - unfamiliar tight shoaling zone; UL - unfamiliar loose shoaling zone.

Shoaling was measured as the time that the subject spent within 10 cm of either shoal (ca. 4 body lengths, as in Magurran et al. 1994 and Griffiths & Magurran 1997a, previous studies documenting familiarity preferences in guppies). Shoaling definitions vary in the literature, with shoaling in small fish is defined as both within 2 or 4 body lengths of other fish (Pitcher & Parrish 1993; Chapman et al. 2008; Swaney et al. 2015; Wright & Krause 2006). We therefore repeated analyses for Tests 1 to 3 with a two-body-length criterion (“tight shoaling”) to check robustness. Results were identical and thus only results using the four-body-length criterion are reported below.

Following a trial, subjects were immediately netted back into their home tanks, adding an opaque white barrier to separate them from stimulus fish of the same tank. No fish was tested more than once. The order of subjects and the position of the bottles was randomized (as in Griffiths & Magurran 1997a) using a random number generator to account for any order or side bias. The water in the tank was mixed using a net between trials to minimize the presence of any localized odour cues and aeration was turned off during trials to avoid uneven dispersal of these cues.

#### Behavioural Measures and Statistical Analysis

All statistical analyses were performed in R, with the R base package (R Core Team, 2022). Effect sizes were calculated with the *effsize* package (Torchiano 2020). Residual diagnostics were performed to assess model fit, including checks for normality (Shapiro-Wilk test) and homoscedasticity of residuals.

We analysed the total time individual guppies from different populations (mixed wild origin and Aripo/Quare-cross) spent with familiar vs. unfamiliar shoals using a linear model, with fish ID and tank ID as fixed effects to account for the repeated structure of data (since each subject could visit both the familiar and unfamiliar sides within the same trial). Experimenters were not blind to experimental conditions, but all behavioural data were extracted using automated video tracking software EthoVision XT v11.567 (Noldus), with manual correction of tracking errors.

### Results and Discussion

All subjects approached at least one of the shoals and spent, on average, 68.2% of the 15-minute test grouping with either shoal in what represents 25.9% of the total tank volume. However, there was no significant difference in the amount of time guppies spent with familiar versus unfamiliar fish (Figure 2; F(1,28) = 0.954, p = 0.34, effect size d = 0.18, 95% CI = - 0.18-0.55), and no significant effect of or interaction with population (Population: F(1,28) = 0.006, p = 0.94; Population*Familiarity: F(1,28) = 0.292, p = 0.59). For exposition we also analysed populations separately, again finding no significant preference for or against familiar fish (mixed wild origin: F(1,14) = 0.163, p = 0.69, effect size d = 0.11, 95% CI = −0.42-0.63; Aripo/Quare: F(1,14) = 0.815, p = 0.38, effect size d = 0.24, 95% CI = −0.29-0.77).

**Figure 2:**
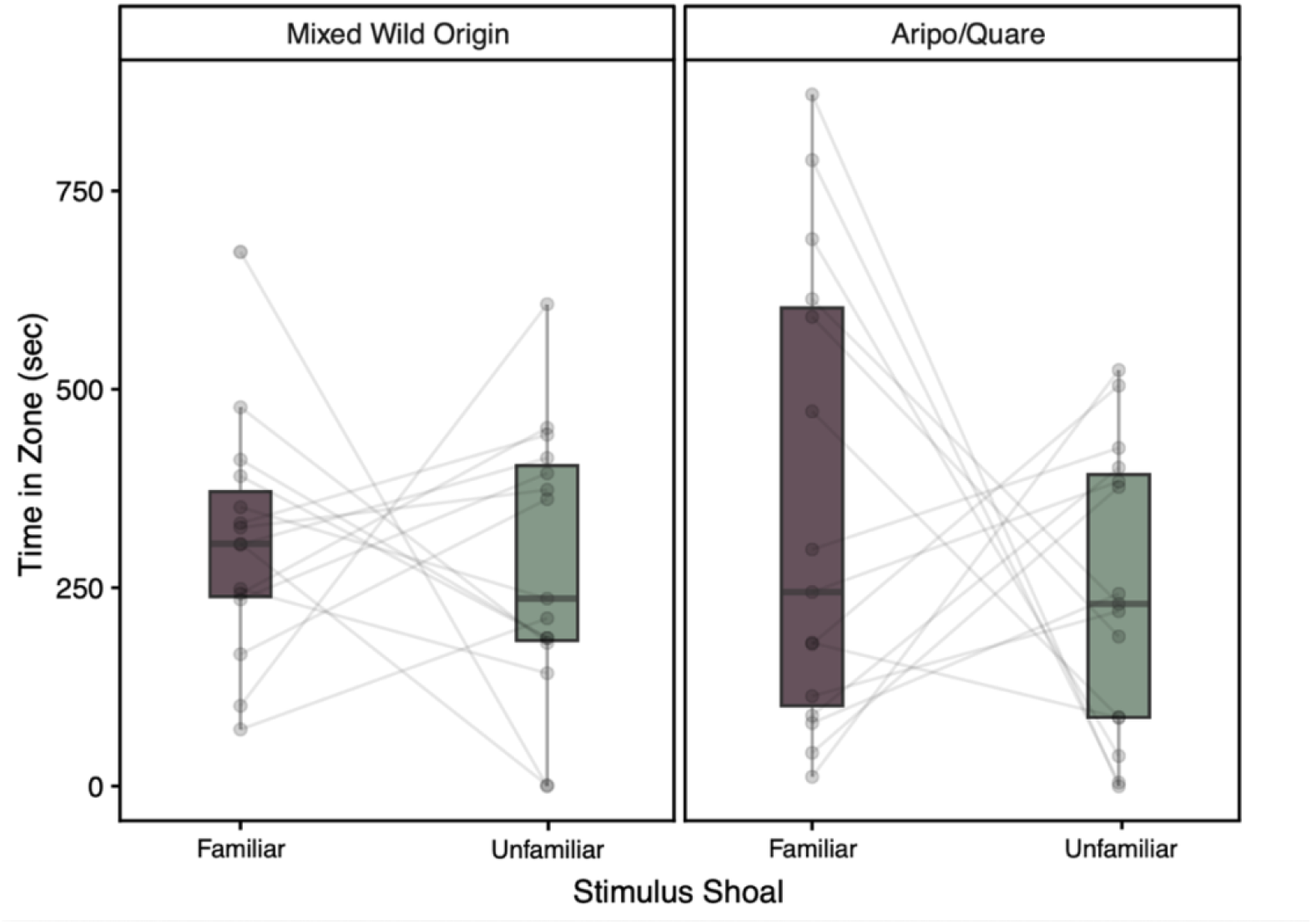
Time spent (in seconds) near an unfamiliar or familiar shoal during Test 1 for a) mixed wild origin and b) Aripo/Quare-cross populations. Points joined by lines represent a single subject (n=30 total, n=15 per population). Boxplots indicate median (horizontal line), first and third quartiles *(box); whiskers extend to the furthest value within 1.5 times the IQR. Populations are plotted separately for exposition but were analysed together*.

We thus saw no indication of a familiarity preference after 15 days of cohabitation. Although our methods and duration of familiarization closely matched previous work that had detected a familiarity preference (Griffiths and Magurran 1997a), we reasoned that perhaps the duration of familiarization or the precise methodology used could have impacted our findings. We therefore retested the same subjects after an additional ∼40 days of familiarization, using a different choice test set-up where the two stimulus shoals were alongside each other instead of on opposite sides of the tank.

## Tests 2 and 3

### Materials and Methods

Set up and housing conditions were generally identical to Test 1, and the same subjects were reused. However, the Aripo/Quare-cross females were housed in pairs to increase the subject sample size and potentially provide increased opportunity to develop a familiarity preference, while the mixed wild origin population was housed in the same groups of six as in Test 1. A different experimenter to Test 1 ran Tests 2 and 3 and fish were not tracked across tests. Fish were housed in these conditions for around 40 days, with tests run over several days, resulting in a total familiarization time of 53-56 days. Two subjects were replaced at the beginning of familiarization for Test 2 (one wild origin and one Aripo/Quare-cross) due to natural deaths, so the two new subjects had a shorter familiarization time than all other subjects (∼40 days).

### Test 2: Familiarity Preference

As in Test 1, individual subjects could choose between a familiar and an unfamiliar stimulus. For wild origin subjects, this was a 3-fish shoal from either the subject’s home tank or a randomly chosen different tank (Figure 3a). For Aripo/Quare-cross subjects, this was an individual fish from either the subject’s home tank or a randomly chosen different tank (Figure 3b). Both stimuli were presented in front of the subject testing tank (*l* 54 x *w* 40 x *h* 40 cm; water at 20 cm), separated from each other by an opaque black partition. Each rectangular shoal zone was 27 cm long. During a 3-minute acclimation period where the subject was placed into a glass cylinder, it could observe both stimuli simultaneously before making a choice. When released following acclimation, behaviour was video recorded for 5 minutes.

**Figure 3:**
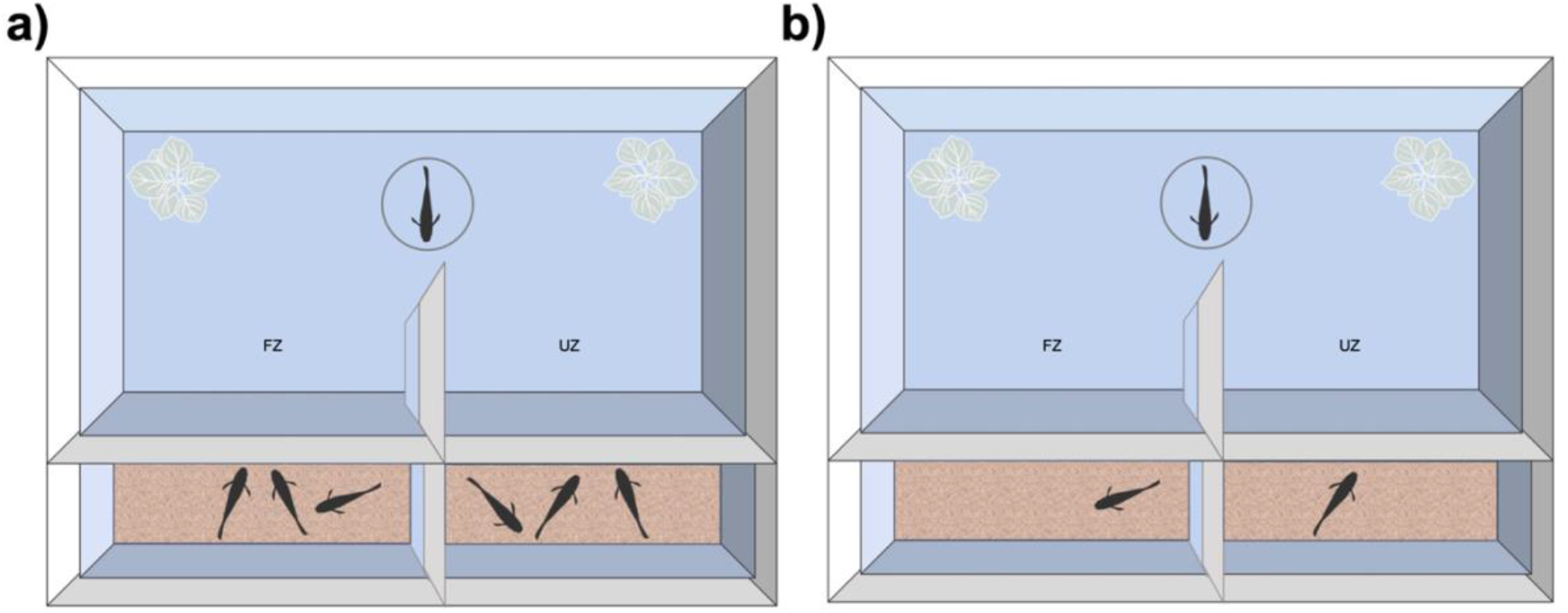
Test 2 familiarity preference test set-up, top view. Subject was released from a central cylinder and allowed to swim freely between familiar and unfamiliar fish in neighbouring tanks. **a) Mixed wild origin population**: subject could group with three familiar or three unfamiliar fish and **b) Aripo/Quare-cross population**: subject could group with one familiar or one unfamiliar fish. Opaque black barriers were placed between the two stimulus tanks and white artificial plants were at either end of the testing tank. The side of the tank with familiar vs. unfamiliar fish was randomly determined. FZ - familiar zone; UZ - unfamiliar zone. Not to scale.

Time shoaling with the familiar and unfamiliar stimuli was measured as in Test 1 (0-5 cm: tight shoaling zone, 0-10 cm: loose shoaling zone). Each test consisted of two trials between which the side of the familiar stimulus was switched to account for any side bias, and the subject was given an additional minute to acclimate. The familiarity preference testing occurred over two days. Subject order and location of unfamiliar stimuli were randomized each day. Unlike Test 1, fish used as stimuli were reused as subjects the following day to increase sample size. To allow identification of these fish, an opaque white barrier was placed between subjects and stimulus fish overnight following the first day of preference testing.

### Test 3: Shoaling Preference

After the familiarity preference test was completed for all subjects, subjects were put back into their familiarity housing tanks and over the following two days we performed a shoaling test with the same individuals. Three Aripo/Quare-cross fish were excluded from testing due to mortality or misidentification.

We had two stimuli - a tank with a social stimulus (a shoal of familiar fish for the mixed wild origin: Figure 4a and a single familiar fish for the Aripo/Quare-cross population: Figure 4b) and an empty tank. Otherwise, the apparatus and methodology were identical to Test 2.

**Figure 4:**
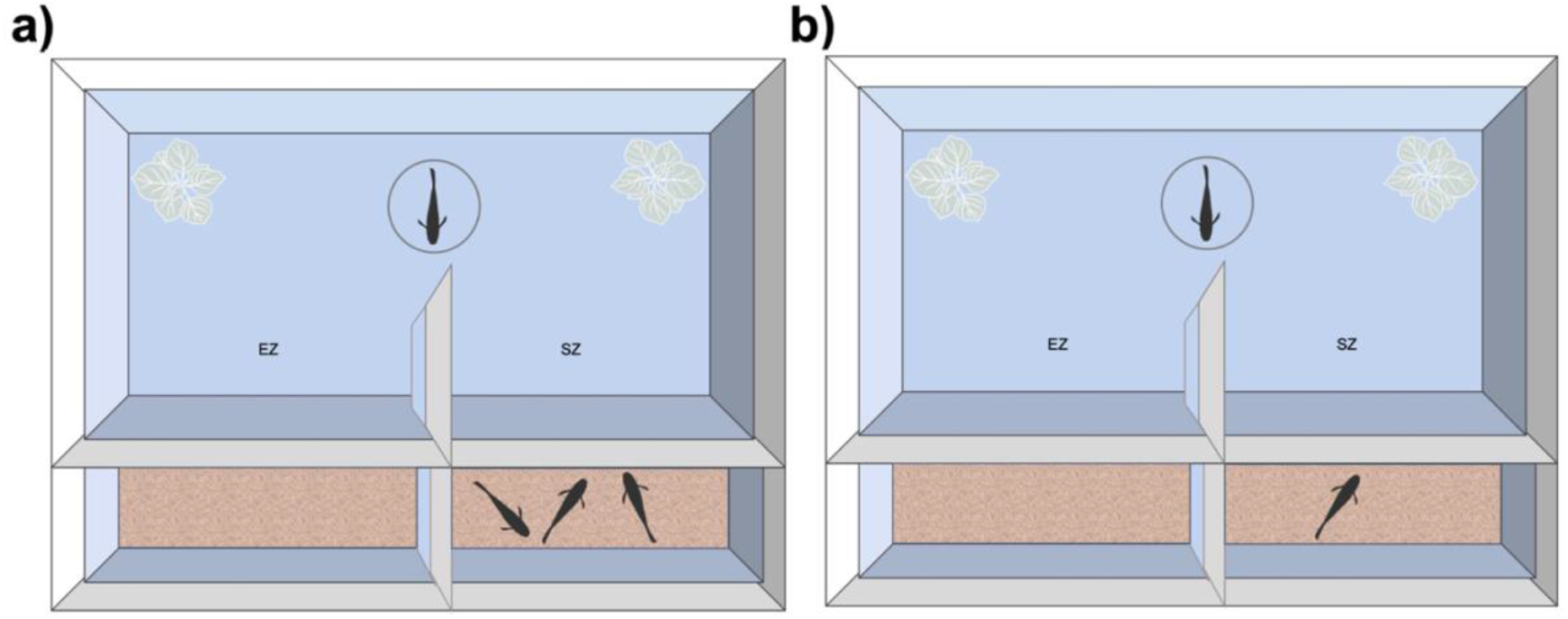
Test 3 shoaling preference test set-up, top view. Subject was released from a central cylinder and allowed to swim freely between a shoal and an empty compartment in neighbouring tanks. **a) Mixed wild origin population**: subject could group with three fish and **b) Aripo/Quare-cross population**: subject could group with one fish. Opaque black barriers were between the two stimulus tanks and white artificial plants were at either end of the testing tank. The side of the tank with the stimulus fish was randomly determined. EZ - empty zone; SZ - shoaling zone. Not to scale.

#### Behavioural Measures and Statistical Analysis

The two populations were analysed separately due to the differences in housing and testing conditions. Time in each zone was averaged across the two trials for analysis. We analysed the total time spent with familiar vs. unfamiliar individuals and the total time spent with the shoal vs. the empty compartment using paired t-tests, since we did not need a model to account for population as was required in Test 1.

## Results

### Test 2: Familiarity preference

There was no significant difference in the time spent with the familiar versus unfamiliar stimulus in both the mixed wild origin (n=25; Figure 5a; paired t-test: t(24) = 0.924, p = 0.36, effect size d = 0.35, 95% CI = −0.43-1.12) and Aripo/Quare-cross populations (n=28; Figure 5b; paired t-test: t(27) = −0.382, p = 0.70, effect size d = −0.11, 95% CI = −0.71-0.49).

**Figure 5:**
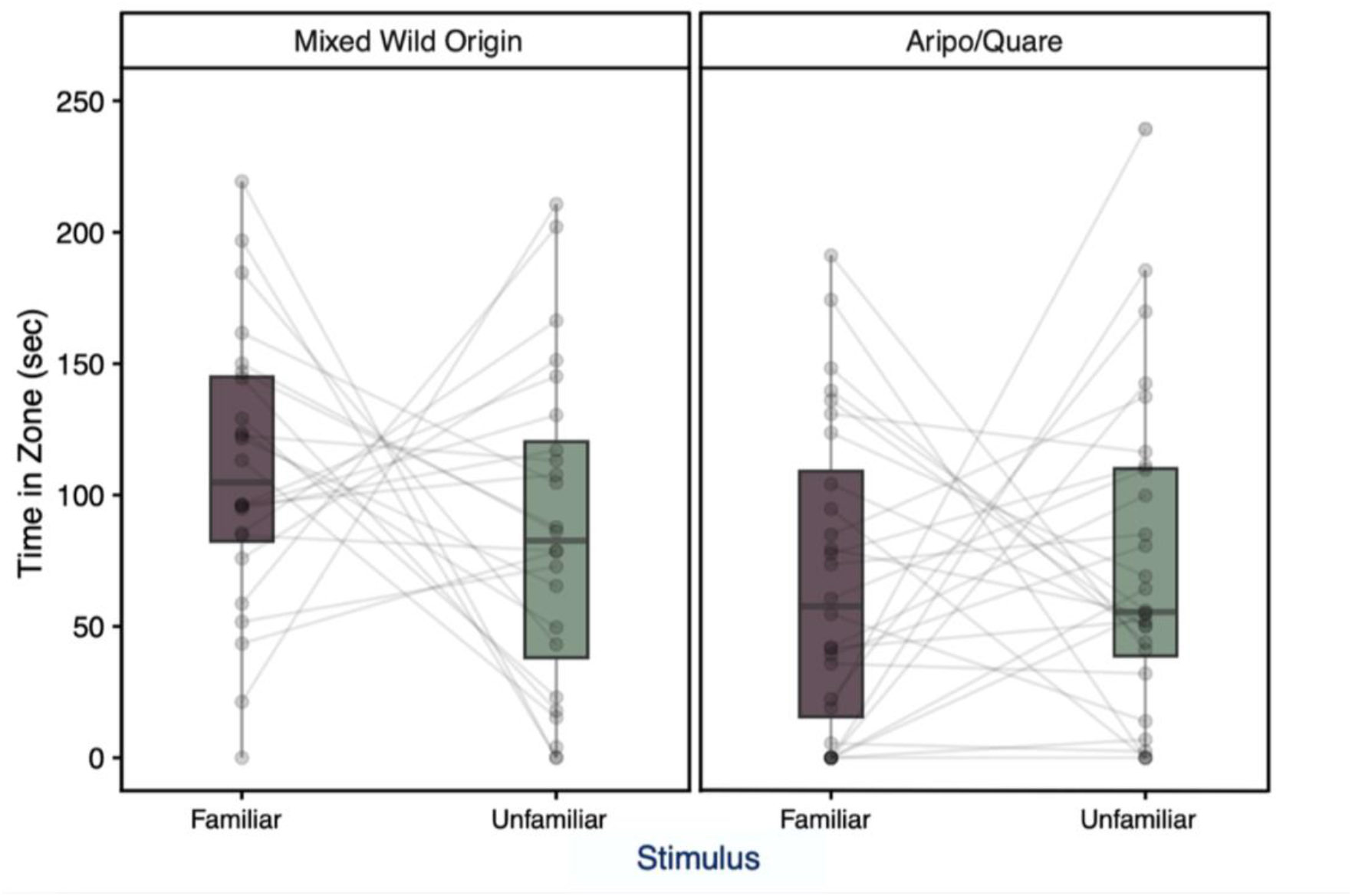
Time spent (in seconds) near unfamiliar or familiar individuals during Test 2 for a) mixed wild origin (n=25) and b) Aripo/Quare-cross (n=28) populations. Points joined by lines represent a single subject. Boxplots indicate median (horizontal line), first and third quartiles (box); whiskers extend to the furthest value within 1.5 times the IQR.

### Test 3: Shoaling preference

Subjects had a significant preference to spend time on the shoal side (versus the empty side) in both the mixed wild origin (n= 25; Figure 6a; paired t-test: t(24) = 6.472, p < 0.0001, effect size d = 2.37, 95% CI = 0.93-3.80) and Aripo/Quare-cross populations (n= 25; Figure 6b; paired t-test: t(24) = 3.612, p = 0.001, effect size d = 1.20, 95% CI = 0.32-2.09).

**Figure 6:**
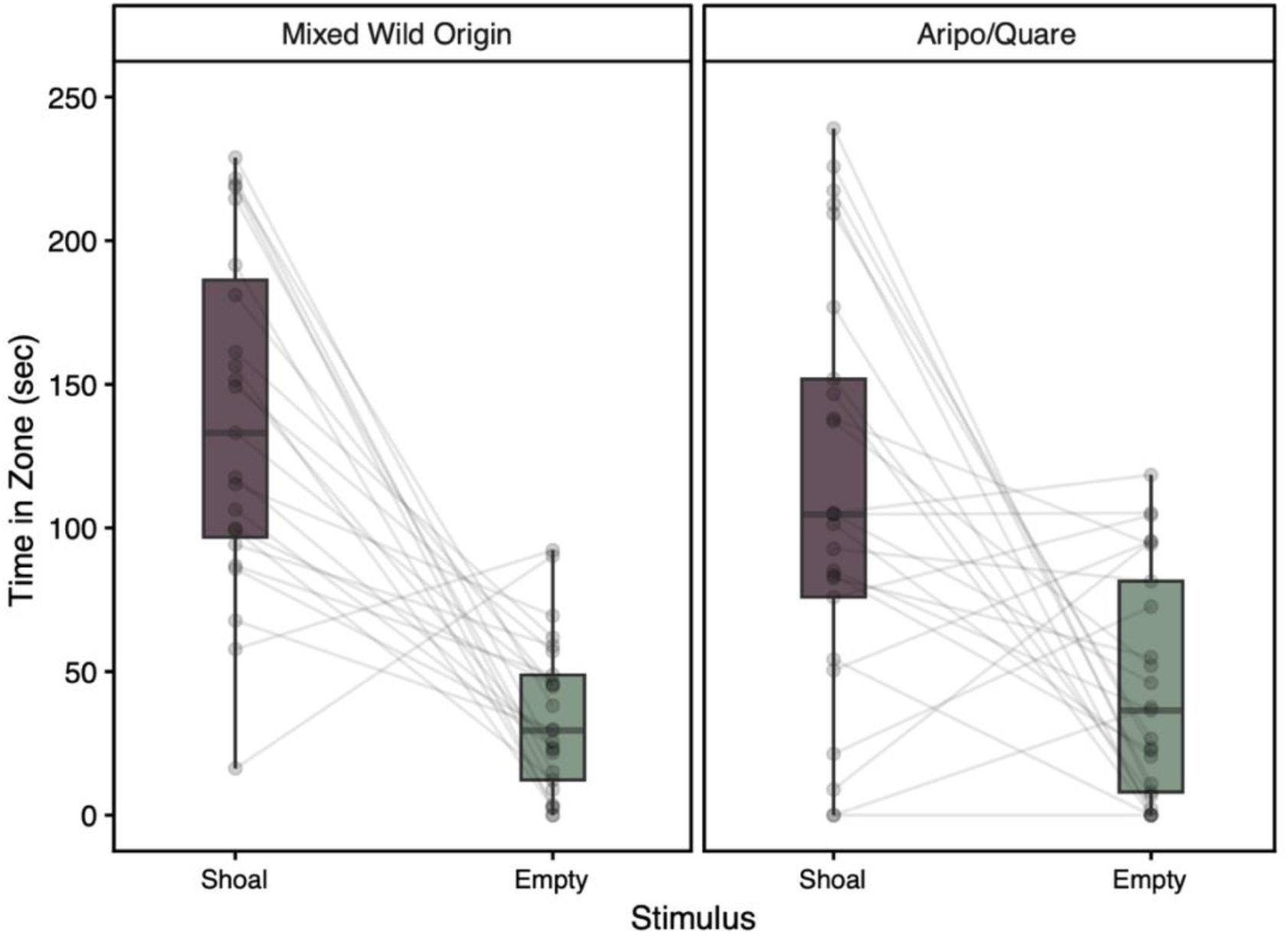
Time spent (in seconds) near other individuals or an empty compartment during Test 3 for a) mixed wild origin (n=25) and b) Aripo/Quare-cross (n=25) populations. Points joined by lines represent a single subject. Boxplots indicate median (horizontal line), first and third quartiles (box); whiskers extend to the furthest value within 1.5 times the IQR.

## General Discussion

Previous research has indicated that female guppies shoal preferentially with tankmates over unfamiliar conspecifics after about 12 days of cohabitation (Griffiths and Magurran 1997a). Our study provided no evidence for a familiarity preference in either of the populations we studied after 15 days of cohabitation using methodology modelled on this previous work, or after 55 days of cohabitation using a modified methodology. However, we did see a strong preference to group with conspecifics (vs. an empty compartment) in our control shoaling test in both populations, as expected (Magurran 2005; Lachlan et al. 1998; Cabrera-Álvarez et al. 2017), suggesting that both populations are able to demonstrate preferences in the experimental apparatus.

Familiarity preferences have been reported in multiple fish species, but with differences in expressed preferences impacted by population, species, sex, context, and other factors (Krause et al. 2000; Griffiths 2003; Frommen et al. 2007; Ward et al. 2009; Griffiths & Ward 2011; Ward et al. 2020). We thus reviewed published work on familiarity preferences in guppies, restricting our examination to familiarity amongst same-sex adults or amongst juveniles (Table 1). We included studies directly testing for familiarity preferences as well as studies examining whether familiarity results in behavioural differences, since the latter indicates that familiarity has developed. We tabulated factors thought to be important influences on the development and expression of a familiarity preference.

**Table 1:** Summary of studies on the effects of same-sex familiarity on guppy (Poecilia reticulata) behaviour. Data are reported on separate rows for different populations, diets, sexes, and the discriminatory cues available (ordered by studies on females, males, and juveniles and by publication date). Three familiarisation methods were used: guppies were housed together, guppies in the same natural pools were considered familiar, or fish in a natural guppy shoal were considered familiar. As noted in the original studies, kinship between fish cannot be excluded as a factor promoting association in natural pools or shoals, while kinship was accounted for by haphazard assignment of fish to groups or by controlling kinship in the group housing studies (e.g. Griffiths & Magurran 1999; Hain & Neff 2007; Hain et al. 2016). Warburton & Lees (1996): we report their results for non-kin. Griffiths & Magurran (1999): familiar and unfamiliar fish were both kin. Hain & Neff (2007): familiarity preferences were not statistically significant for tests within kin and within non-kin, but were significant when these data were pooled. Price & Rodd (2006): familiar fish were also full or half-siblings. Magurran et al. (1994) presented combined results for the two populations but noted no significant population differences in shoaling time. TT: Trinidad and Tobago (native populations), HP: high-predation regime (see Magurran 2005), AUS: Australia (feral populations).

| Author(s) | Year | Familarisation Method | Test | Cues Available | Source Population | Subject Origin | Diet | Subject Sex | Stimulus Sex | Kinship accounted | Expressed Preference or Behaviour |
|---|---|---|---|---|---|---|---|---|---|---|---|
| Magurran et al. | 1994 | Group housing | Two-choice shoaling test | Visual | Lower Aripo TT | Captive-bred | Tetramin flake food | Females | Females | Yes | Familiar |
| Magurran et al. | 1994 | Group housing | Two-choice shoaling test | Visual | Lower Turure TT | Captive-bred | Tetramin flake food | Females | Females | Yes | Familiar |
| Griffiths and Magurran | 1997a | Group housing | Two-choice shoaling test | Visual and chemosensory | Lower Tacarigua TT | Captive-bred | Tetramin flake food | Females | Females | Yes | Familiar |
| Griffiths and Magurran | 1997b | Natural pools with <40 females | Nearest neighbour | All (free-swimming) | Upper Tunapuna TT | Wild-born | Wild | Females | Females | No | Familiar |
| Griffiths and Magurran | 1997b | Natural pools with >60 females | Nearest neighbour | All (free-swimming) | Upper Tunapuna TT | Wild-born | Wild | Females | Females | No | None |
| Griffiths and Magurran | 1998 | Natural shoals | Two-choice test | Visual and chemosensory | Tacarigua HP TT | Wild-born | Wild then Tetramin flake food | Females | Females | No | Familiar |
| Lachlan et al. | 1998 | Group housing | Two-choice test | Visual | Domestics | Domestic | Flake food | Females | Females | Yes | Familiar |
| Swaney et al. | 2001 | Group housing | Foraging task | All (free-swimming) | Domestics | Domestic | Flake food | Females | Females | Yes | Faster feeding with familiar demonstrators |
| Godin et al. | 2003 | Natural pools | Two-choice shoaling test | Visual and chemosensory | Upper Tunapuna TT | Wild-born | Wild then NutraFin flake food | Females | Females | No | None |
| Bhat and Magurran | 2006 | Group housing | Two-choice shoaling test, shelter emergence | Visual and chemosensory | Lower Tacarigua TT | Captive-bred | *Artemia franciscana* | Females | Females | Yes | Familiar, faster shelter emergence with familiars |
| Morrell et al. | 2007 | Group housing | Two-choice shoaling test | Visual and chemosensory | Domestics | Domestic | Bloodworm (*Chironomus* spp.) | Females | Females | Yes | Familiar |
| Morrell et al. | 2007 | Group housing | Two-choice shoaling test | Visual and chemosensory | Domestics | Domestic | Flake food | Females | Females | Yes | Unfamiliar |
| Morrell et al. | 2008 | Natural shoals | Shoaling, foraging task | All (free-swimming) | Arima TT | Wild-born | Wild then flake food | Females | Females | No | Faster feeding, smaller groups and lower association strength in natural shoals |
| Ward et al. | 2009 | Group housing | Two-choice shoaling test | Visual and chemosensory | Tacarigua TT | Captive-bred | Bloodworm or *Daphnia* | Females | Females | Yes | Familiar |
| Sievers and Magurran | 2011 | Group housing | Two-choice shoaling test | Visual and chemosensory | Lower Tacarigua TT | Captive-bred | Tetramin flake food | Females | Females | Yes | Familiar |
| Barbosa et al. | 2016 | Group housing | Shoaling, novel area entrance | All (free-swimming) | Lower Tacarigua TT | Captive-bred | Flake food | Females | Females | Yes | Familiar (shoaling), Unfamiliar (novel area entrance) |
| Barbosa et al. | 2016 | Group housing | Shoaling, novel area entrance | All (free-swimming) | Pitch Lake TT | Captive-bred | Flake food | Females | Females | Yes | Familiar (shoaling), Unfamiliar (novel area entrance) |
| Davis et al. | 2017 | Group housing | Group polarization | All (free-swimming) | Darwin AUS | Wild-born | Wardley tropical flake food | Females | Females | Yes | Familiar |
| Hasenjager and Dugatkin | 2017 | Group housing | Fission–fusion behavior | All (free-swimming) | Quare TT | Captive-bred | Tetramin flake food | Females | Females | Yes | None |
| Cattelan et al. | 2019 | Group housing | Two-choice shoaling test | Visual and chemosensory | Tacarigua TT | Captive-bred | *Artemia salina* nauplii and DuplarinS granulated food | Females | Females | Yes | None, though individual consistency in familarity preferences |
| Crane et al. | 2020 | Group housing | Sensitivity to disturbance cues | Chemosensory | Lopinot TT | Wild-born | Wild then flake food | Females | Females | Yes | Significant response only to cues from groups of familiarised individuals |
| Crane et al. | 2020 | Group housing | Sensitivity to disturbance cues | Chemosensory | Upper Aripo TT | Wild-born | Wild then flake food | Females | Females | Yes | Significant response to cues regardless of group familiarisation |
| Santacà et al. | 2021 | Group housing | Two-choice shoaling test | Conflicting visual and chemosensory | Lower Tacarigua TT | Captive-bred | Aqua tropical flake food | Females | Females | Yes | None, though initially spent more time in the compartment with the familiar visual stimuli |
| De Meester et al. | 2025 | Group housing | Two-choice shoaling test | Visual | Large-brained selection lines | Captive-bred selection lines | Flake food and *Artemia* sp. | Females | Females | Yes | Familiar |
| De Meester et al. | 2025 | Group housing | Two-choice shoaling test | Visual | Small-brained selection lines | Captive-bred selection lines | Flake food and *Artemia* sp. | Females | Females | Yes | Familiar |
| Present study |  | Group housing | Two-choice shoaling tests | Visual and chemosensory | Lower Aripo/Quare cross TT | Captive-bred | TetraMin flake food and *Artemia sp.* | Females | Females | Yes | None |
| Present study |  | Group housing | Two-choice shoaling tests | Visual and chemosensory | Mixed wild TT origin | Captive-bred | TetraMin flake food and *Artemia sp.* | Females | Females | Yes | None |
| Present study |  | Group housing | Two-choice shoaling tests | Visual | Lower Aripo/Quare cross TT | Captive-bred | TetraMin flake food and *Artemia sp.* | Females | Females | Yes | None |
| Present study |  | Group housing | Two-choice shoaling tests | Visual | Mixed wild TT origin | Captive-bred | TetraMin flake food and *Artemia sp.* | Females | Females | Yes | None |
| Griffiths and Magurran | 1998 | Natural shoals | Two-choice shoaling test | Visual and chemosensory | Tacarigua HP TT | Wild-born | Wild then Tetramin flake food | Males | Males | No | None |
| Godin et al. | 2003 | Natural pools | Two-choice shoaling test | Visual and chemosensory | Upper Tunapuna TT | Wild-born | Wild then NutraFin flake food | Males | Males | No | None |
| Croft et al. | 2004 | Group housing | Two-choice shoaling test | Visual and chemosensory | Arima TT | Wild-born | Champion flake food | Males | Males | Yes | Familiar |
| Price and Rodd | 2006 | Group housing | Male-male interactions | All (free-swimming) | Quare TT | Captive-bred | Flake food or live brine shrimp nauplii. | Males | Males | No | More aggression between unfamiliar fish |
| Granroth-Wilding and Magurran | 2013 | Group housing | Shelter emergence, shoaling, food competition, mating competition | All (free-swimming) | Lower Tacarigua TT | Captive-bred | Tetramin flake food | Males | Males | Yes | Familiar (following, shoaling, food competition); None (mating competition) |
| Barbosa et al. | 2016 | Group housing | Shoaling, novel area entrance | All (free-swimming) | Lower Tacarigua TT | Captive-bred | Flake food | Males | Males | Yes | None (shoaling), Unfamiliar (novel area entrance) |
| Barbosa et al. | 2016 | Group housing | Shoaling, novel area entrance | All (free-swimming) | Pitch Lake TT | Captive-bred | Flake food | Males | Males | Yes | None (shoaling), Unfamiliar (novel area entrance) |
| Sogawa et al. | 2023 | Pair housing | Sequential aggression test | Visual | Neon tuxedo domestics | Domestics | Tetramin flake food | Males | Males | Yes | Familiar |
| De Meester et al. | 2025 | Group housing | Two-choice shoaling test | Visual | Large-brained selection lines | Captive-bred selection lines | Flake food and *Artemia* sp. | Males | Males | Yes | Familiar |
| De Meester et al. | 2025 | Group housing | Two-choice shoaling test | Visual | Small-brained selection lines | Captive-bred selection lines | Flake food and *Artemia* sp. | Males | Males | Yes | Familiar |
| Warburton and Lees | 1996 | Group housing | Two-choice shoaling test | Visual | Domestics | Captive-bred | Tetra Guppy Food Special Diet | Juveniles | Juveniles | Yes | Familiar (one measure only) |
| Warburton and Lees | 1996 | Group housing | Two-choice shoaling test | Mainly chemosensory | Domestics | Captive-bred | Tetra Guppy Food Special Diet | Juveniles | Juveniles | Yes | None |
| Griffiths and Magurran | 1999 | Group housing | Two-choice shoaling test | Visual | Lower Tacarigua TT | Captive-bred | Tetramin flake food | Juveniles | Juveniles | Yes | Familiar |
| Griffiths and Magurran | 1999 | Group housing | Two-choice shoaling test | Chemosensory | Lower Tacarigua TT | Captive-bred | Tetramin flake food | Juveniles | Juveniles | Yes | Familiar |
| Hain and Neff | 2007 | Group housing | Two-choice shoaling test | Visual and chemosensory | Paria TT | Captive-bred | Not stated | Juveniles | Juveniles | Yes | Familiar |
| Hain et al. | 2016 | Group housing | Two-choice shoaling test | Visual and chemosensory | Lower Oropouche TT | Captive-bred | Brine shrimp, Tetra flake food | Juveniles | Juveniles | Yes | None |
| Hain et al. | 2016 | Group housing | Two-choice shoaling test | Visual and chemosensory | Tunapuna TT | Captive-bred | Brine shrimp, Tetra flake food | Juveniles | Juveniles | Yes | None |
| Hain et al. | 2016 | Group housing | Two-choice shoaling test | Visual and chemosensory | Upper Yarra TT | Captive-bred | Brine shrimp, Tetra flake food | Juveniles | Juveniles | Yes | None |
| Hain et al. | 2016 | Group housing | Two-choice shoaling test | Visual and chemosensory | Upper Aripo TT | Captive-bred | Brine shrimp, Tetra flake food | Juveniles | Juveniles | Yes | Familiar |
| Hain et al. | 2016 | Group housing | Two-choice shoaling test | Visual and chemosensory | Lower Guanapo TT | Captive-bred | Brine shrimp, Tetra flake food | Juveniles | Juveniles | Yes | None |
| Hain et al. | 2016 | Group housing | Two-choice shoaling test | Visual and chemosensory | Paria TT | Captive-bred | Brine shrimp, Tetra flake food | Juveniles | Juveniles | Yes | Familiar |
| Wolcott et al. | 2017 | Group housing | Escape response | All (free-swimming) | Lower Tacarigua TT | Captive-bred | Tetramin flake food | Juveniles | Juveniles | Yes | Lower responsiveness in familiar groups, no effects on escape latency or magnitude |

While numerous studies find evidence for familiarity effects in guppies of different origins (wild, wild-type raised in captivity, feral, and domestic), both in the field and captivity, this was not universal and variation is observed both between and within populations (Table 1). It is possible that additional negative findings may be unpublished (‘the file-drawer problem’; Rosenthal 1979; Sánchez-Tójar et al. 2018; Howard & Barron 2024). However, numerous factors can influence the outcome of animal studies, including factors not usually the focus of biological study, such as characteristics of the laboratory or experimenter (Webster & Rutz 2020; von Kortzfleisch et al. 2022). Ecological, evolutionary, contextual, and methodological factors could all impact the development and expression of familiarity preferences. For example, in guppies, variation in familiarity preferences were found across six wild populations that were laboratory-reared and tested under the same conditions, suggesting evolved differences (Hain et al. 2016; Table 1). Godin et al. (2003) suggested within-population differences in familiarity preferences could be due to recent environmental conditions in the wild, while Morrell et al. (2007) found housing diet could alter familiarity preferences. Sex differences in familiarity preferences are also often described (Table 1). Our review did not clearly identify a single factor as responsible for our results, and we suspect their most likely explanation is a combination of population, rearing and methodological effects.

Work with zebrafish has found evidence for familiarity preference only when shoals were 30 cm apart, not at 45 or 60 cm (Swaney et al. 2025), raising the possibility that what might appear to be minor methodological details can affect outcomes. Our Test 1 set-up was very close to that of Griffiths and Magurran (1997a), with both using 1 litre perforated containers for the shoals to allow both visual and chemosensory cues, shoals separated by 62 cm, 20 cm water depth, a 10 cm shoaling zone and a 15 minute trial. We did use a smaller stimulus shoal (we used 3 fish as in Bhat and Magurran (2006) instead of 4), more days of cohabitation (15 days instead of 12), a larger sample, and different guppy populations. While Griffiths and Magurran (1997a) began seeing a familiarity preference on day 12, it peaked around day 15, which is why we chose this for Test 1. Morrell et al. (2007) found that female guppies fed flake food preferred unfamiliar fish, while other studies feeding either flake food alone or flake food and *Artemia sp*. as in our study found preferences for familiar fish (Table 1). Familiarity preferences have also been described for guppies bred in captivity. We used two mixed-origin populations, one based on a cross of two populations from high-predation regimes in Trinidad, another of mixed Trinidadian origin. Guppy populations vary considerably in behaviour (Magurran 2005); it is possible that population origin could explain our results, but we cannot exclude other factors such as rearing or laboratory conditions, or interactive effects. Guigueno et al. (2025) compared two guppy populations bred in common-garden conditions, finding effects of a stressor on social learning depending on the population, indicating interactive effects of environmental factors and genetic origin. While we cannot firmly identify the reason our results differ from similar studies, negative results are important in establishing patterns and we hope building additional understanding of familiarity.

It is well appreciated that methodological details, genetic background, and rearing differences can have large impacts on the ability to measure behaviours and performance and importantly, to compare them between studies (Stewart et al. 2012; O’Neill et al. 2018; Webster and Rutz 2020; Jones et al. 2023). For instance, Stewart et al. (2012) found that fish scale their locomotory activity corresponding to the size of the tank, and O’Neill et al. (2018) demonstrated that acclimation time can affect behavioural repeatability. In light of this, nuanced investigations of how specific discrepancies affect the detection and measurement of subject participation and behaviour are a recommended next step (Salena et al. 2021; Jones et al. 2023).

Associating with familiar conspecifics has been found to increase the benefits of grouping in fish, leading to more cohesive shoals, effective information transfer, and faster learning (Chivers et al. 1995; Barber and Wright 2001; Swaney et al. 2001; Lucon-Xiccato et al. 2022). Even though multiple past studies have seen a preference form in guppies (Table 1), we did not find similar evidence in our current study. This suggests that familiarity might be contextual in nature, only developing or expressed under specific conditions or in certain populations. Further research could investigate how context, population, and set-up can play a role in familiarity preference and its development over time. Specifically, formal comparisons of populations and environment would provide valuable information on the specific conditions under which familiarity preferences develop both in the laboratory and in the wild. Similarly, investigating methodological aspects including shoal size, distance of the stimulus shoals from the subject, and acclimation time are all valuable next steps. Studies like this one contribute to the growing knowledge on the conditions under which grouping preferences are most beneficial.

## Acknowledgements

We thank Elinor Lynch for assistance with body length measurements and Adam Crane and Gilles De Meester for confirming methodological details.

## Ethics Statement

The work was approved by the McGill University Animal Care Committee (protocol MCGL-7708), and followed applicable laws and ASAB/ABS guidelines.

## Data Availability Statement

The data that supports the findings of this study are available at https://doi.org/10.5683/SP3/LVQKZ2.

## CRediT Author Contributions

ASR: conceptualization, data curation, formal analysis, investigation, methodology, visualization, writing – original draft, writing – review and editing; NPCA: investigation, methodology; JTS: conceptualization, funding acquisition, supervision, writing – review and editing; SMR: conceptualization, funding acquisition, resources, supervision, writing – review and editing.

